# The effect of the Icelandic mutation APP^A673T^ in the line 66 model of tauopathy

**DOI:** 10.1101/2025.10.09.681356

**Authors:** Anne Anschuetz, Lianne Robinson, Miguel Mondesir, Valeria Melis, Bettina Platt, Charles R. Harrington, Gernot Riedel, Karima Schwab

**Author notes:** Corresponding author: Gernot Riedel. Authors with equal contribution (Gernot Riedel, Karima Schwab).

## Abstract

The Icelandic mutation in the amyloid precursor protein (APP), APP^A673T^, has been identified in Icelandic and Scandinavian populations and is associated with a significantly lower risk of developing Alzheimer’s disease (AD). Although this mutation led to reduction in amyloid β-protein (Aβ) production, its effect on tau pathology is not well studied. We have crossed line 66 (L66) tau transgenic mice that overexpress the P301S aggregation-prone form of tau with C57Bl6/J mice expressing a single point mutation edited into the murine APP gene via CRISPR-Cas gene editing, termed APP^A673T^. We have performed ELISA, histopathological and behavioural analyses of heterozygous male/female L66 and L66xAPP^A673T^ crosses at the age of 6 months to investigate the effect of the A673T mutation on tau brain pathology and behavioural deficits in these mice. Using immunohistochemistry, we found only a moderate, yet significant, reduction of mAb 7/51-reactive tau in prefrontal cortex for L66xAPP^A673T^ compared to L66 mice. Quantification of tau in soluble/insoluble brain homogenate fractions by ELISA confirmed the lack of overt differences between genotypes, as did our extensive behavioural phenotyping using six different paradigms accessing motor function, olfaction, depression/apathy-like behaviour, as well as exploration and sociability. Therefore, the APP^A673T^ mutation does not appear to modulate tau pathology or motor and neuropsychiatric behaviour in L66 tau transgenic mice.

## Introduction

Dementia, such as Alzheimer’s disease (AD), is an incurable disease characterised by a progressive, and irreversible decline of cognition leading to disoriented behaviour, altered personality, aggression, agitation, anxiety, difficulties with speech and comprehension, and impaired gait and movement (Scott & Barrett, 2007). The decline is related to a loss of neuronal function caused by the formation of neurofibrillary tangles and extracellular, neuritic plaques(Alzheimer, 1907). These two hallmarks of AD consist mainly of the microtubule-associated protein tau (Goedert et al., 1988; Wischik et al., 1985; Wischik et al., 1988; Wischik et al., 1988; Wischik & Crowtber, 1986) with amyloid β-protein (Aβ) present also in neuritic and non-neuritic plaques (Eanes & Glenner, 1968; Glenner & Wong, 1984; Glenner & Wong, 1984; Masters et al., 1985). Tau protein was identified as a polymerisation factor for microtubules, which promotes microtubule assembly and provides axonal stabilisation (Weingarten et al., 1975). In addition, tau may also be involved in the regulation of axonal guided transport through interactions with motor proteins and other binding partners (Mietelska-Porowska et al., 2014). Amyloid-β is generated through the cleavage of the amyloid precursor protein (APP), a protein that functions in a variety of physiological processes, including modulation of synaptic function, facilitation of neuronal growth and survival, and protection against oxidative stress (Bishop & Robinson, 2004; Coulson et al., 2000).

Early work has shown that tau deletion of tau leads to altered microtubule organisation and complex motor deficits including impaired performance on the rotarod and decreased locomotion in the open field test (Harada et al., 1994; Ikegami et al., 2000), while deletion of the amyloid precursor protein has also been associated with hypoactivity and reduced grip strength (Zheng et al., 1995). By contrast, overexpression of mutated tau in mice led to its accumulation in nerve cell bodies, axons and dendrites causing muscle atrophy and dysfunctional sensorimotor performance (Götz et al., 1995; Probst et al., 2000; Spittaels et al., 1999), while more physiological expression led to phenotypes akin to FTD without motor impairments (Koss et al., 2016). Similarly, some transgenic APP mice also developed neuritic plaques, synaptic loss, and memory impairments (Dodart et al., 1999; Games et al., 1995; Sturchler-Pierrat et al., 1997). While the loss of normal function and gain of toxic function of tau and Aβ are clearly implicated in the pathophysiology of AD, for other dementias like frontotemporal dementia (FTD) these pathways do not necessarily interact (Citron et al., 1994; D’Souza et al., 1999; Giaccone et al., 2010; Goate et al., 1991; Goedert, 2005; Yoshida et al., 2002). In cultured hippocampal neurons, however, degeneration required the presence of both tau and Aβ-treatment (Rapoport et al., 2002), and that reducing endogenous tau levels without altering Aβ prevented behavioural deficits in Aβ-transgenic mice, emphasising that tau may be essential for Aβ-induced neurotoxicity (Roberson et al., 2007). On the other hand, human tau increased Aβ levels in Aβ-transgenic mice and reducing tau simultaneously lowered Aβ plaque formation (Bright et al., 2015).

To further address the interaction between Aβ and tau in the pathogenesis of AD, FTD, and related disorders we have crossed APP^A673T^ mice, expressing murine Aβ with the Icelandic mutation, with line 66 (L66) tau transgenic mice. L66 mice overexpress the P301S aggregation-prone form of tau and are characterised by early-onset tau pathology, and subsequent sensorimotor and motor dysfunction (Lemke et al., 2020; Melis et al., 2015; Schwab et al., 2021, 2025). In humans, the P301S mutation is associated with FTD with Parkinsonism (Spillantini et al., 1998). APP^A673T^ mice harbour the Icelandic mutation in the murine APP gene generated by CRISPR-Cas gene editing technology. This mutation, first identified in elderly Icelandic and later in different Scandinavian populations, significantly reduces risk of developing AD in carriers (Jonsson et al., 2012; Martiskainen et al., 2017). The protective effect of A673T is believed to be primarily achieved through reduced β-cleavage of APP (Blennow et al., 2006; Hampel et al., 2021; Xia et al., 2021). The A673T mutation proved protective against Aβ pathology in Aβ-transgenic mice (Shimohama et al., 2024) but its effect on tau pathology/aggregation in the absence of Aβ overexpression remains unknown. In this exploratory study, heterozygous L66xAPP^A673T^ crosses were analysed both behaviourally and histopathologically relative to L66 tau transgenic mice. Multiple secondary endpoints included determination of synaptic proteins and behavioural phenotypes. We assumed that, if protective, the Icelandic mutation should reduce tau levels in L66xAPP^A673T^ relative to L66 mice but failed to observe any such effect.

## Materials and Methods

### Study design

The study was exploratory. The primary read-out was tau pathology and whether it is reduced in L66xAPP^A673T^ compared to L66. No power calculations were performed a priori but the group sample size was based on experience and expectations from previous experiments performed on L66 mice (Lemke et al., 2020; Melis et al., 2015; Robinson et al., 2024; Schwab et al., 2021). The body weight of mice was determined prior to experimental start and additionally once per week during the experimentation period. Welfare recommendations suggested any mouse with a body weight decline ≥ 15% to be excluded from the study but this did not occur, and no mouse was excluded during the *in vivo* phase of the study. A randomisation sequence for behavioural testing was generated using the random function in Excel (Microsoft Office) to generate 2 cohorts balanced for sex and genotype. Experimenters and care takers were blinded to the genotype of the mice during group allocation, behavioural assessment, data acquisition and primary data analyses. A different investigator performed statistical analyses for behavioural data and was not blinded to any study details. Following tissue collection, an independent experimenter, also blinded to the genotype of mice, performed immunohistochemistry, ELISA assays, and all statistical analyses relating to these measurements.

### Animals

All animal experiments were performed in accordance with the European Communities Council Directive (63/2010/EU) with local ethical approval under the UK Animals (Scientific Procedures) Act (1986) and its amended regulations (2012), and under the project licence number PP2213334 compliant with the ARRIVE guidelines 2.0 (Percie du Sert et al., 2020).

Mice were bred commercially in positive-pressure isolators (Charles River, Margate, UK). Homozygous tau transgenic line 66 mice, expressing the full-length human tau transgene htau40 (amino acids 1-441) with two mutations in position 301 (P301S) and 335 (G335D) (Melis et al., 2015). Mice were bred on a white NMRI background strain. Mice harbouring the Icelandic mutation were generated by CRISPR-Cas gene editing technology with a nucleotide modification of the murine APP gene, changing amino acid 673 from alanine to threonine, on a black C57Bl6/J background at Jax laboratories (Bar Harbor, USA), termed APP^A673T^ mice. Here, crosses were bred from homozygous L66 (male or female) with heterozygous APP^A673T^ (male or female) mice, and the resulting agouti-coloured offsprings were all heterozygous for the L66 htau40 transgene, but ear biopsies were genotyped for the A673T mutation in the APP gene by Transnetyx Inc. (Cordova, USA). These were either A673T positive or wild type. A total of fifty-nine mice, 6-month-old, were included in the study (Table 1). Experimental mice were delivered by truck to the animal facilities at the University of Aberdeen, Scotland, one month prior to the start of the experiments. Mice were kept in sex-specific litters ≥2 in stock box open housing under constant environmental conditions (20-22°C temperature, 50-65% humidity, an air exchange rate of 17-20 changes per hour, and a 12-h light/dark cycle with lights turned on at 7 am with simulated sunrise/sunset) and ad libitum chow (Special Diet Services, Witham, UK) and water throughout. Mice were also provided with corncob bedding, paper strips, and cardboard tubes (DBM Scotland Ltd, UK) as enrichment throughout the experiment. One week before and throughout behavioural testing, all animals were singly housed to harmonise holding conditions throughout the in-live phase of the experiment. Behavioural testing commenced with motor coordination on the rotarod followed by home cage observations in the PhenoTyper, nest building in the home cage, sucrose preference test, buried cookie test and social interaction (for details, see below). They were maintained singly housed in Macrolon type III cages in the same holding room except when they were transferred to specialised testing rooms for behavioural assessments or for euthanasia and sacrifice. Mice were given ample time to recover between experiments (>3 days) and 30 minutes of acclimatisation to behavioural rooms before any procedure took place.

**Table 1:** Study groups and cohort sizes. L66: heterozygous line 66 mice, L66xAPP^A673T^: crosses carrying both the L66 tau-transgene and the A673T mutation in the APP gene. N: number of mice.

|  | N Male (cohort 1 & cohort 2) | N Female (cohort 1 & cohort 2) |
| --- | --- | --- |
| L66 | 14 (7 & 7) | 14 (7 & 7) |
| L66xA673T | 14 (7 & 7) | 17 (9 & 8) |
| N - Total | $\Sigma$ 59 | |

### Behavioural testing

Behavioural testing was conducted using a battery of tests which consisted of assessments of motoric (Rotarod), exploratory (home cage activity analysis in the PhenoTyper), social interaction, buried cookie test, and apathy-like/anhedonic behaviours (nest building and sucrose preference tests) performed over four weeks. The age of 6 months was chosen because at this age L66 mice have developed extensive tau pathology in the brain that leads to behavioural dysfunction such as in the rotarod and the nest building test (Lemke et al., 2020; Melis et al., 2015; Robinson et al., 2024; Schwab et al., 2021).

#### Rotarod

To access motor function, a four-lane rotarod system was used (Model 33700-R/A; TSE Technical & Scientific Equipment GmbH, Bad Homburg, Germany). Animals were tested for 4 trials per day over 3 consecutive days with inter trial intervals of 2 – 3 minutes during which the apparatus was cleaned with 70% ethanol. Four mice were run simultaneously, with the run sequence randomised and counterbalanced for time of testing and rod location using a Latin square design. At the start of a trial the mouse was placed on a slowly rotating rod (1 rpm (rotation per minute)) with its head facing against the rotation. The trial was started once all animals were in position and the rotation speed of the rod was accelerated from 1 rpm to 45 rpm over a period of 5 minutes. The trial was ended when the animal fell off the rod or when the maximum trial time was reached. The time spent on the rod (sec) for each of the 12 trials (T1-T12) was extracted as the primary read-out and used for analyses.

#### PhenoTyper

Following the rotarod test, locomotor activity and circadian activity patterns were monitored using the PhenoTyper home cage observation system (Noldus Information Technology, Wageningen, The Netherlands). The cages consisted of clear Perspex walls and opaque floors (30 x 30 x 35 cm) and contained a food hopper and water bottle allowing free access to food and water during testing. The cages were filled with sawdust and the top unit of each cage contained a built-in digital infrared sensitive video camera and infrared lighting sources for video tracking. The infrared sources enabled continuous behavioural recordings in both dark and light periods. Animals were individually placed in the PhenoTyper cages with the environmental conditions identical to those of the holding rooms. The animals were housed in PhenoTypers for 7 consecutive days and were given two days of habituation followed by assessment of circadian activity from days 3 −7. The behaviour of the animals was recorded using the video-tracking software EthoVision XT (Version 16, Noldus Information Technology) by background subtraction at 25Hz. Habituation to the novel environment was measured by analysing the distance moved during the initial 3 hr period following placement in the PhenoTyper with data extracted in 10-minute time bins. During the experimental days, the distance travelled was extracted and analysed in 1 hr time bins to determine the exploratory and circadian activity of the mice. Thirty mice were run simultaneously.

#### Nest building test

During single housing in Macrolon Type III cages (Tecniplast, Milan, Italy), a cotton nestlet (50 mm x 50 mm square pressed cotton, DBM Scotland Ltd, UK) was positioned in the cage prior to the start of the dark cycle and the nest building ability of the mice was visually scored after a period of 16 hr (Day 1) and 48hrs (Day 2). The scoring was performed by three independent researchers blind to the genotype of the mice using the scoring system developed by Deacon (Deacon, 2012) and recently validated by Robinson et al (Robinson et al., 2024). The scoring system utilised a 5-point scale determined by the completeness of the nest. A score of 1 was assigned if the nestlet remained predominantly untouched. A partially torn up nestlet was given a score of 2 whilst an almost entirely shredded nestlet but no clear nest location scored 3. Once the nestlet was entirely shredded and a nest area evident a score of 4 was assigned for a flat nest with a maximum score of 5 only given when the nest resembled a crater with walls higher than the height of the mouse. The scores of the 3 researchers were averaged for each mouse and used for statistical analysis.

#### Sucrose preference

For the sucrose preference test, animals were housed in Perspex cages (54 ×50 x 37 cm) (Ugo Basile, Comero, Italy). Each cage was filled with corn cob bedding and equipped with two drinking bottles and a food hopper. Two days following nest building, the animals were individually placed into sucrose test cages for two days of habituation. Subsequently, water in one bottle was exchanged for 1% sucrose solution and the weights of the two bottles recorded. The position of the sucrose bottle was counterbalanced (left or right) for animals/groups. Both bottles were weighed after 24 hrs and the position of the sucrose bottle alternated to avoid any spatial preference. The weights of the two bottles were recorded again following a further 24 hrs, after which the animals were returned to their home cages. Water and sucrose consumption for each animal was averaged for the 48-hr period and sucrose preference determined as the amount of sucrose solution consumed divided by the total intake of fluid multiplied by 100. To control for possible water/sucrose leakages from the bottles small bags were attached to the bottle holders close to the spouts and used to collect any fluid which was then subtracted from the bottle weights. Upon completion of the 48-hour test, animals were returned to their Macrolon home cages.

#### Buried cookie test

Mice were initially habituated to a piece of cookie (McVities Hobnobs) introduced into their home cages on alternate days, at least 5 days before the test commenced. Prior to the test animals were food-restricted overnight (up to 16 hrs) although free access to water was maintained throughout this period. The test was conducted in Macrolon Type III cages (Tecniplast, Milan, Italy) measuring 36.9 x 16.5 x 13.2 cm illuminated by indirect lighting (∼60 lux). The flooring of the cages was filled with corn cob bedding to a depth of 2 cm and a piece of cookie was buried approximately 0.5 cm below the surface of the bedding. A transparent plastic lid with holes was positioned on top of the cage during testing to prevent the animal from climbing out. Animals were subjected to 3 trials with a maximum trial duration of 5 minutes per trial and an inter-trial interval of 1 minute. Trial 1 was a habituation session without any cookie. Whilst during trials 2 and 3 a piece of cookie was buried, with the position of the cookie switched and fully counterbalanced between trials to prevent any spatial bias. At the start of each trial the animal was placed in the centre of the cage and allowed to explore freely with the behaviour monitored by an overhead video camera and locomotor activity tracked by ANY-maze (version 5.1, Stoelting Co, Dublin, Ireland). The main parameter recorded was the latency for the animal to place both forepaws on the cookie piece during the trial duration; this was measured both manually by the experimenter using a stopwatch and automatically by the tracking software. A latency of 300 seconds was recorded if the subject failed to find the cookie within the allotted time. After completion of all trials, animals were returned to their home cages and given food immediately. The mean latency of trials 2 and 3 was calculated for each animal and used for analysis.

#### Social interaction

Social interaction and recognition of the mice was recorded using a three-chambered arena (Platt et al., 2011; Riedel et al., 2009). The arena was constructed of white Perspex (63 cm x 42 cm x 22 cm) and consisted of two outer chambers (each 22 cm x 42 cm) and a central compartment (19 cm x 42 cm) separated by dividers with apertures allowing free movement between the chambers. A cylinder with metal bars was positioned in the centre of each of the outer chambers and used for the confinement of a stranger mouse during social interaction. The apparatus was illuminated by indirect overhead lighting during testing (∼60 lux). Test sessions consisted of three phases: i) habituation, ii) sociability and iii) social memory/recognition. During habituation, the animal was placed in the central compartment and allowed to freely explore the arena for 10 mins. In the sociability phase an unfamiliar mouse (stranger 1 (S1)) of the same sex as the test mouse was placed inside one of the social interaction cylinders in either the left or right chamber whilst the cylinder in the opposite chamber remained empty. The location of the stranger was randomly selected and counterbalanced (left or right) across groups. The test animal was again released in the central compartment and allowed to freely explore the arena over a 10-min test session. Measures taken included time spent within the immediate vicinity (4 cm interaction zone) of the two cylinders (S1 or empty). During the social memory test a novel stranger was introduced into the second, previously empty cylinder and the exploration of the two cylinder zones (novel, unfamiliar stranger (S2) and now familiar stranger (S1)) were recorded for a further 10 minutes. The inter-trial interval between the different phases was 5 mins and between animals the arena and cylinders were all cleaned and disinfected with 70% ethanol. Activity and exploratory behaviour of the mice were video-recorded and stored online (ANY-maze). Social interactions were quantified as time spent with S1 compared to the corresponding empty compartment (sociability), or with S2 compared to S1 (social memory). Distance travelled, time spent interacting (total), the time spent interacting with a stranger, and the discrimination index (ratio of time spent interacting with a stranger mouse to the total time spent interacting) were extracted/calculated and used for statistical analyses.

### Tissue collection

Brain tissue was harvested from all fifty-nine mice that underwent behavioural testing. Mice were euthanised via intraperitoneal injections of a sub-lethal dose of sodium pentobarbital (Dolethal (200 mg/ml), Covetrus, UK). Mice underwent intra-cardiac perfusion with heparinised phosphate-buffered saline (0.1 M PBS with 0.05% (v/w) heparin, pH 7.4) for 5 minutes. Skulls were then removed and the whole brain immediately dissected out. The right brain hemisphere was separated, fixed overnight at room temperature in 10% (v/v) neutral-buffered formalin, dehydrated and embedded in paraffin. Sagittal sections were prepared at 5 µm using a rotary microtome (HM 325, Leica Biosystems, Sheffield UK), and mounted onto glass slides (SuperFrost^TM^, Thermo Fisher Scientific, Lutterworth UK). Sections were collected from regions at interaural 0.96 to 1.44 mm lateral of midline (Paxinos & Franklin, 2019), and three sections were collected on one slide for each mouse and antibody. The left-brain hemisphere was transferred immediately to liquid nitrogen after brain removal and kept at −80°C until used for protein extraction/ELISA protein quantification.

### Immunohistochemistry (IHC) and quantification of tau

Sagittal sections were stained in a sex specific way using two immunohistochemistry staining boxes for male and two for female samples. Each box included a balanced number of L66 and L66xAPP^A673T^ mouse brains. Sections (three sections per mouse and antibody) were stained according to our standard protocol (Lemke et al., 2020) using three different antibodies: the mouse monoclonal antibody mAb 7/51 targeting the epitope 350-368 within the microtubule domain of the tau protein (TauRx Therapeutics, Singapore, diluted 1: 1,000); the mouse monoclonal antibody HT7 targeting the epitope 159-163 within proline-rich domain of tau (Thermo Scientific, Loughborough UK, #MN1000, diluted 1: 1,000); the mouse monoclonal antibody mAb 27/499 targeting the N-terminal epitope 14-26 (TauRx Therapeutics, diluted 1: 200). All chemicals were purchased from Merck Millipore (Burlington, MA, USA). Images of cornu ammonis (CA1), the dentate gyrus (DG), the visual cortex (CTX), the prefrontal cortex (PFC), and the cerebellum (CB) were taken using a light microscope at a 100x magnification (Axio Imager M1, Carl Zeiss, Jena, Germany) and saved in TIFF file format. Microscopic images were analysed using ilastik (Version 1.4.0.post1, (Berg et al., 2019)) and Fiji (Version 2.14.0, (Schindelin et al., 2012)). The pixel classification tool in ilastik enabled training of the software based on a small subset of samples and then apply them to larger sets of images (Berg et al., 2019). Models were trained to segment images into positively stained pixels and non-stained background tissue or artefacts. After applying models to all images, the percentage of positively stained area of the whole image was quantified using Fiji. This was repeated for each of the three antibodies from all fifty-nine mice.

### Protein extraction

All chemicals were purchased from Merck Millipore (Burlington, MA, USA) if not otherwise stated. The left hemibrains were pulverized in a liquid nitrogen prechilled stainless steel mortar and pestle (BioPulverizer, BioSpec, Oklahoma, USA) and homogenized with a pestle and hammer. RIPA lysis buffer (Thermo Fisher Scientific, #89900) containing Pierce Protease and Phosphatase Inhibitor Mini Tablets (Thermo Fisher Scientific, # A32959) was added in a ratio of 5:1 (mL buffer to mg wet tissue) and the homogenate incubated for 30 minutes on ice with casual agitation. After centrifugation at 19,000 g for 2 hours at 4 °C (Centrifuge 5427 R – Microcentrifuge, Eppendorf, Stevenage UK, using the FA-45-48-11 rotor), the supernatant (referred to as the RIPA-soluble supernatant fraction S1) was transferred into new reaction tubes. The residual pellet was homogenised in 5 volumes tris buffered saline (pH 7.6) containing 5 M guanidine hydrochloride (GuHCl) and incubated with mild agitation (11 rotations per minute, Multi Bio RS-24, Biosan, Riga, Latvia) for 16 hours at room temperature. After centrifugation at 15,000 g for 30 minutes at room temperature, the resultant GuHCl supernatant fraction (referred to as S2) was transferred into new tubes. RIPA-soluble (S1) and RIPA-insoluble (S2) fractions were stored at −20 °C until use. Total protein concentration of S1 and S2 fractions were determined by bicinchoninic acid (BCA) protein assay (Pierce™ BCA Protein Assay Kit, Thermo Fisher Scientific, #23225) using bovine serum albumin (BSA) (0.125 – 2.000 mg/ml) as a reference standard.

### Quantification of tau, Aβ and synaptic proteins using ELISA

RIPA-soluble S1 was used to measure non-phosphorylated tau (non-pTau, ROBOSCREEN #847-0108000102), Aβ40 (Invitrogen, #KMB3481), Aβ42 (Invitrogen, #KMB3441), mouse synaptosomal associated protein 25kDa (SNAP25, MyBiosource, San Diego USA, #MBS451917), mouse syntaxin 1A (STX1A, MyBiosource #MBS452386), and mouse synaptophysin (SYP, MyBiosource #MBS453910). For non-pTau quantification, S1 samples were diluted 1:5,000 in assay buffer. For Aβ40 and Aβ42, S1 samples were used undiluted. For synaptic protein ELISA S1 samples were first diluted to a protein concentration of 4µg/µl in RIPA and then further diluted in PBS, as recommended by manufacturer, at a dilution of 1:2 for STX1A and 1:10 for SYP and SNAP25. Undiluted GuHCl S2 samples were used to assess aggregated tau (ROBOSCREEN #847-0104000116). S2 samples were diluted 1:10 to measure Aβ40 and Aβ42 (same kits as above). ELISA assays were performed as per the manufacturer’s instructions. The levels for non-pTau, aggregated tau, SNAP25, SYP and SNTX1A from all fifty-nine mice were analysed. For Aβ ELISA some S1 and S2 samples had low sample volumes and could therefore not be included; where this is the case sample sizes were amended in the figure legends.

### Data analysis

No a priori exclusion criteria were set. However, some IHC samples were excluded due to tissue damage during sectioning or lack of staining possibly due to sample preparation error. Details are specified in the respective sections. All other data are included in the behavioural and cellular analysis and are presented here. Behavioural data were assessed for normality using the Shapiro-Wilk test and Gaussian distribution assumptions were met. Nest building scores were analysed using Aligned Rank Transform (ART) analysis of variance (ANOVA) with repeated measures and time, sex and genotype as independent variables (Kay, 2020). All other data were analysed using either 2-way ANOVA with genotype and time/trial, or sex and genotype as independent variables or 3-way ANOVA with genotype, sex and time as independent variables. GraphPad Prism software (version 10.2.3; GraphPad Software Inc., Boston, USA) was used to generate graphs and conduct statistical analyses. IHC and ELISA data was analysed and graphs generated in R (Version 4.4.3, R Core Team, Vienna, Austria) using linear models and 2-way ANOVA. Where appropriate, *post-hoc* tests were performed using Bonferroni corrections. For IHC staining, males and females were analysed separately and the effects of brain region, genotype and their interaction on percentage area stained were assessed. For each analysis, it was first determined whether data met assumptions for 2-way ANOVA (normality of residuals, heteroscedasticity) or whether data transformations were required. Data transformations used were square root, log or Box-Cox transformation and details are given in the respective figure legends. Data met necessary assumptions after transformation. A similar approach was taken for ELISA read-outs, where the effect of sex, genotype and their interaction on protein levels were analysed. Due to the large number of samples multiple ELISAs were performed, which in part were from different lots and performed on different days. This was accounted for by including a nuisance factor in the analysis for aggregated tau and synaptic proteins. All statistical outcomes are reported based on linear models of transformed data, but figures show untransformed raw data. For each genotype and sex Pearson correlation matrices were generated from ELISA and IHC data and compared visually and statistically using the Jennrich test (Jennrich RI, 1970)to determine if the matrices were significantly different from each other. Extremely high values were removed from ELISA data to avoid skewing results and obtain correlations not driven by potential outliers. IHC data was averaged across brain regions for this analysis. All data are presented as Mean +/− Standard Deviation (S.D.) and alpha was set to p < 0.05.

## Results

We have used L66 mice that overexpress the P301S aggregation-prone form of human tau and crossed them with mice carrying the A673T Icelandic mutation in the murine APP gene that is protective against Aβ aggregation. The experiments performed in this study were aimed at examining the effects of the Icelandic mutation on i) tau and Aβ levels, ii) synaptic protein abundance, and iii) motor and neuropsychiatric behaviour in L66 and L66xAPP^A673T^ mice.

All mice were in good health with no obvious negative impact of tau overexpression or APP-A673T expression at the age of 6 months when they were investigated. The body weights did not differ significantly between genotypes (F_Genotype_(1,55) = 1.00, *p* = 0.32) in male (L66: 39.5±4.8 g vs. L66xAPP^A673T^: 37.8±3.8) or female mice (L66: 32.3±2.7 g vs. L66xAPP^A673T^: 32.1±3.5). However, male mice were heavier than female mice (F_Sex_(1,55) = 42.44, *p* < 0.0001), and this was the case in both genotypes (F_Interaction_ < 1).

### Icelandic mutation and tau pathology

The levels of non-phosphorylated tau (non-pTau) and aggregated tau were measured in the RIPA-soluble and -insoluble fractions, respectively, of whole brain homogenates using ELISA (Fig.1). Levels of non-pTau were similar in L66 and L66xAPP^A673T^ male mice and showed little variability (Fig.1A). While female L66xAPP^A673T^ mice also showed similar levels, female L66 had slightly higher levels and greater variability than the other three groups. Statistical analyses using two-way ANOVA did not return any significances (all F values < 1). The levels of aggregated tau in the RIPA-insoluble fraction were also similar across both genotypes and sexes, with average levels between 5.9-6.2pg/mL for all groups (Fig.1B). Again, neither sex nor genotype or their interaction influenced aggregated tau levels (F values < 1).

**Figure 1:**
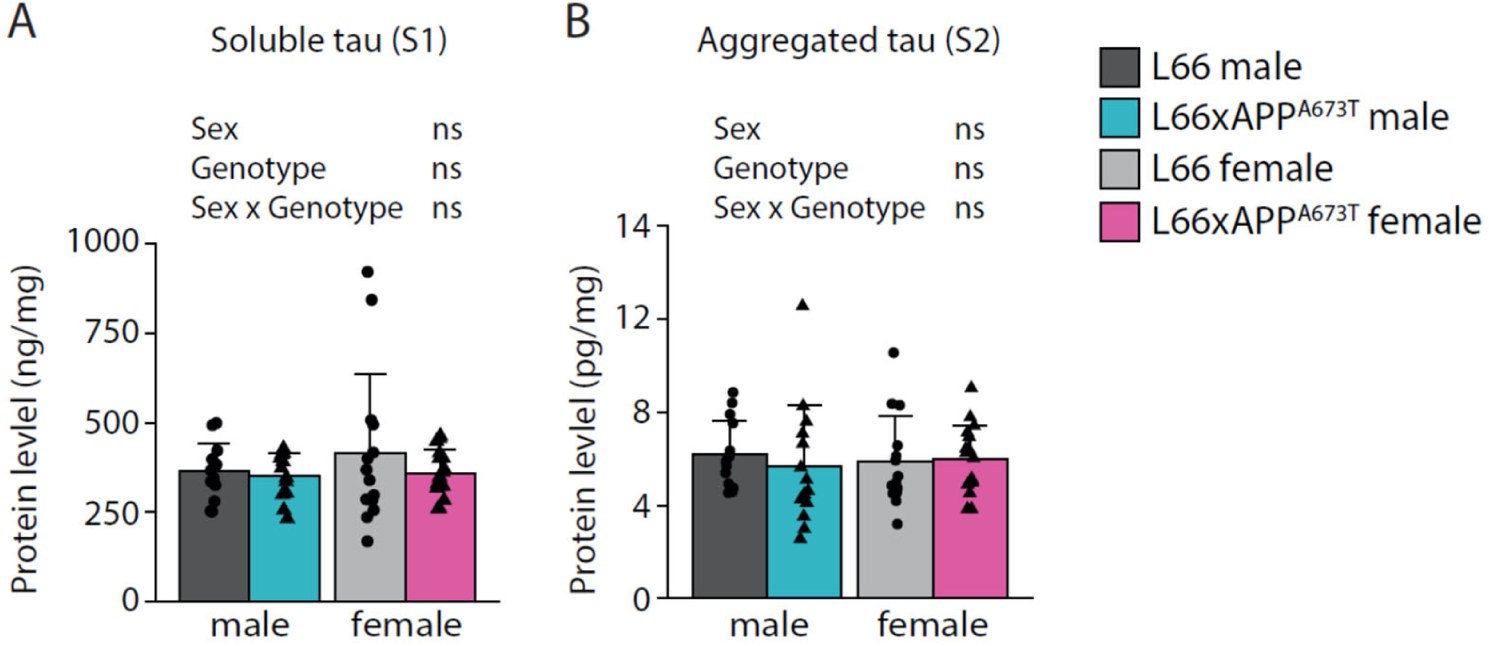
Soluble and aggregated tau ELISA. (A) Total non-phosphorylated tau was quantified in RIPA-soluble S1 fractions. (B) Aggregated tau was quantified in RIPA-insoluble (=GuHCl-soluble) S2 fractions. S1 and S2 fractions were prepared from whole brain homogenates. Raw data for tau was normalised to protein levels and is shown as individual values, with group mean and S.D. Data were analysed using 2-way ANOVA with sex and genotype as independent variables and an additional nuisance factor (batch) for aggregated tau. Data were Box-Cox transformed for analysis. L66: males=14, females =14; L66xAPP^A673T^: males=14, females=17. Abbreviation: ns: not significant.

To analyse tau on a brain-region level, immunohistochemistry was performed using antibodies that target different epitopes of tau, and the positive signal was quantified as percentage of stained area in five regions of interest: CA1, DG, CTX, PFC and CB (Figs.2-4).

The phosphorylation-independent mAb 7/51 binds to residues 350-368 within the microtubule binding domain and the core fragment of tau in the paired helical filament (PHF) typical for AD brain tissue. Positive staining with 7/51 showed a similar pattern in both L66 and L66xAPP^A673T^ crosses, with prominent staining in soma of pyramidal cells in CA1, DG hilus and CTX as well as in large Purkinje cells of the CB (Fig.2A, black arrowheads). A less prominent dendritic staining was also observed in hippocampal and visual and prefrontal cortical areas (Fig.2A, white arrowheads). In male mice (Fig.2B), there was regional variation of mAb 7/51-positivity with greatest levels being observed in cortical areas (F_Brain region_(4,111) = 72.22, *p* < 0.001). Reactivity was similar in L66 and L66xAPP^A673T^ mice (F_Genotype_ < 1), with no obvious interaction between genotype and region factors (F_Interaction_(4,111) = 1.14, *p* = 0.34).

**Figure 2:**
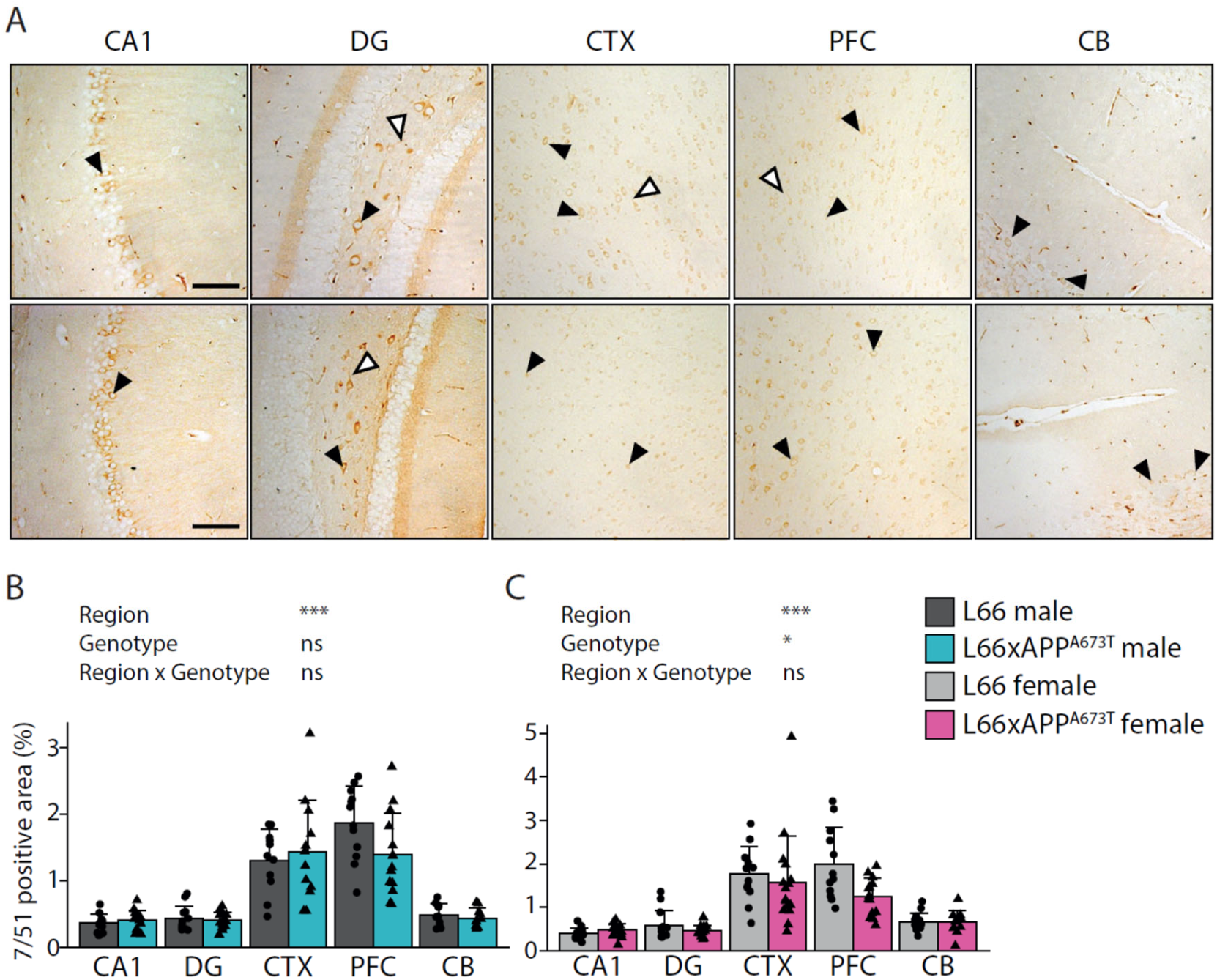
Tau immunohistochemistry using the microtubule domain mAb 7/51. (A) Representative tau immunohistochemistry images in brains of L66 (top) and L66xAPP^A673T^ (bottom) mice stained with the monoclonal anti-tau mAb 7/51. Images from CA1, DG, CTX, PFC, and the CB were taken using a light microscope at a 100x magnification. *Black arrowheads* = cytosolic staining; *white arrowheads =* axonal/dendritic staining; *scale bars*, 100µm. Tau levels were quantified as stained area in percent using ilastik and Fiji and is shown for male (B) and female (C) mice as individual values, with group mean and S.D. Data were analysed using 2-way ANOVA with genotype and region as independent variables (* p < 0.05; *** p < 0.001). Data was log transformed for analysis. Males: L66: n=12 (DG n=11, CB n=10); L66xAPP^A673T^: n=14 (CTX n=12, CB n=10). Females: L66: n=14 (CTX n=13, PFC n=12); L66xAPP^A673T^: n=16 (PFC n=14, CB n=12). Abbreviations: CA1: hippocampal CA1, CB: cerebellum, CTX: visual cortex, DG: dentate gyrus, ns: not significant, PFC: prefrontal cortex.

Nonetheless, a trend towards decreased in 7/51-reactivity was observed in PFC. In female mice (Fig.2C), again staining intensity was highest in cortical regions (F_Brain region_(4,131) = 61.15, *p* < 0.001). Importantly, female L66xAPP^A673T^ crosses exhibited less 7/51-reactive tau than L66 females (F_Genotype_(1,131) = 4.47, *p* = 0.036), and this difference was most notable in CTX and PFC.

HT7 is a further phosphorylation-independent anti-tau antibody which recognises an epitope in the proline-rich region of the tau protein, a region involved in the interaction of tau with actin and other cytoskeletal proteins. Intraneuronal staining in CA1, hilus and granular cell layer of the DG as well as visual and prefrontal cortex were observed; staining patterns were similar in L66 and L66xAPP^A673T^ crosses (Fig.3A, black arrowheads). Processes were also frequently labelled in CA1, but less prominently in DG and cortical regions (Fig.3A, white arrowheads). The CB was devoid of any labelling. The staining intensity with HT7 was greatest in CTX and PFC, both in male (F_Brain region_(3,95) = 52.33 *p* < 0.001, Fig.3B) and female mice (F_Brain region_(3,114) = 66.99, *p* < 0.001, Fig.3C), confirming the staining pattern already seen with mAb 7/51 (see above). Furthermore, no differences were observed between L66 and L66xAPP^A673T^ crosses, either in male or female cohorts (all F values < 1).

**Figure 3:**
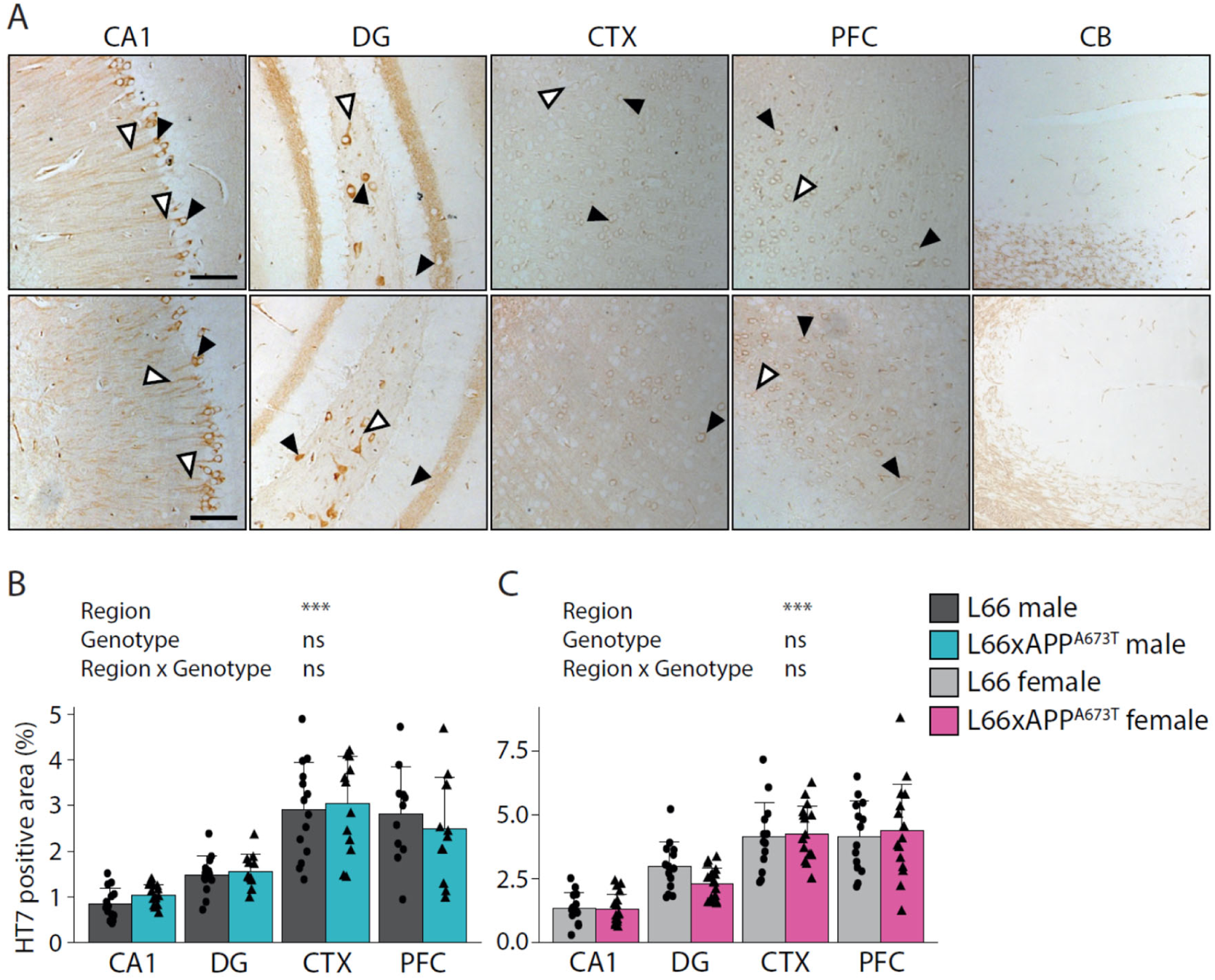
Tau immunohistochemistry using the proline-rich domain antibody HT7. (A) Representative tau immunohistochemistry images in brains of male and female L66 and L66xAPP^A673T^ mice stained with the monoclonal anti-tau antibody HT7. Images from CA1, DG, CTX, PFC, and the CB were taken using a light microscope at a 100x magnification. *Black arrowheads =* cytosolic staining; *white arrowheads =* axonal/dendritic staining; *scale bars*, 100µm. Tau levels were quantified as stained area using ilastik and Fiji and are shown for male (B) and female (C) mice as individual values, with group mean and S.D. Data were analysed using 2-way ANOVA with genotype and region as independent variables (*** p < 0.001). Data was log transformed for analysis. Males: L66: n= 14 (PFC n=11); L66xAPP^A673T^: n=14 (DG n=11, CTX n=13, PFC n=12). Females: L66: n=14; L66xAPP^A673T^ n=17 (PFC, CTX n=16). Abbreviations: CA1: hippocampal CA1, CB: cerebellum, CTX: visual cortex, DG: dentate gyrus, ns: not significant, PFC: prefrontal cortex.

The N-terminal, phosphorylation-independent anti-tau antibody 27/499 binds the epitope of tau known to interact with cytoplasmic components/proteins. The staining pattern with this antibody resembled the pattern revealed by the two antibodies described above, although the intensity was slightly weaker overall and completely absent in CB (Fig.4A). Additionally, the staining did not differ across brain regions, either in males (Fig.4B,) or females (Fig.4C). Like the HT7 labelling reported above, no differences for 27/499-reactive tau were observed between L66 and L66xAPP^A673T^crosses in male or female cohorts.

**Figure 4:**
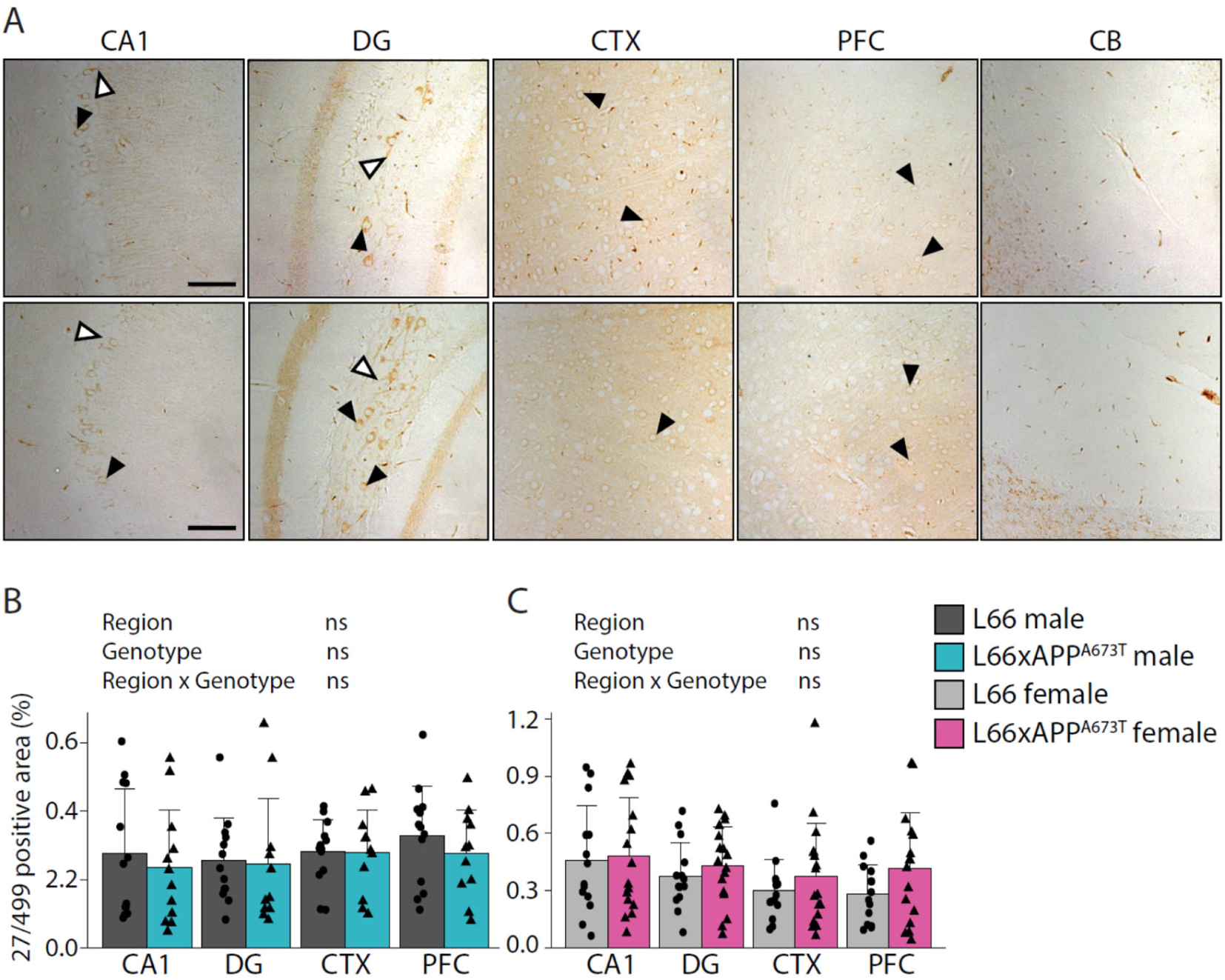
Tau immunohistochemistry using the N-terminal domain antibody 27/499. (A) Representative tau immunohistochemistry images in brains of male and female L66 and L66xAPP^A673T^ mice stained with the monoclonal anti-tau antibody 27/499. Images from CA1, DG, CTX, PFC, and the CB were taken using a light microscope at a 100x. *Black arrowheads* = cytosolic staining; *white arrowheads =* axonal/dendritic staining; *scale bars*, 100µm. Tau levels were quantified as stained area using ilastik and Fiji and are presented for male (B) and female (C) mice as individual values, with group mean, and S.D. Data were analysed using 2-way ANOVA with genotype and region as independent variables. No data transformation was required. Males: L66: n=13; L66xAPP^A673T^: n=12, (DG, CTX, PFC n=11). Females: L66: n=14; L66xAPP^A673T^: n=17 (CA1, PFC n=16). Abbreviations: CA1: hippocampal CA1, CB: cerebellum, CTX: visual cortex, DG: dentate gyrus, ns: not significant, PFC: prefrontal cortex.

### Icelandic mutation, Aβ and synaptic proteins

To assess effects of the protective Icelandic mutation on Aβ production in L66 and L66xAPP^A673T^ crosses, Aβ40 and Aβ42 were analysed in S1 and S2 fractions. Aβ40 in S1 fractions was lower in L66xAPP^A673T^ compared to L66 (F_Genotype_(1,55): 9.51, *p* = 0.003; Fig.5A), while Aβ42 and Aβ42/Aβ40 ratios in the same fraction were similar in both sexes and genotypes (Fig.5B and 5C). Furthermore, no differences were observed between genotypes for the RIPA-insoluble Aβ40, Aβ42 and Aβ42/Aβ40 ratio in S2 fractions (Fig.5D-F). Interestingly, Aβ40 in S1 was greater in female mice compared to male mice in both genotypes (F_Sex_(1,55) = 8.32, *p* = 0.006; Fig.5A). Having established a small, yet significant, reduction of tau and soluble Aβ40 in L66xAPP^A673T^, we further explored whether this reduction may lead to changes in synaptic protein expression using a set of three presynaptic proteins, SYP, SNAP25 and STX1A that were quantified by ELISA. The levels of SYP showed a high degree of variability but were overall similar across genotypes and sexes (F values < 1; Fig.5G). SNAP25 had slightly higher abundance than SYP (SNAP25: 8.85±1.89ng/mg vs. SYP: 7.18±3.52 ng/mg) and showed lower variability (Fig.5H). Again, there were no significant differences in SNAP25 levels between L66 and L66xAPP^A673T^, and the levels were also comparable between male and female cohorts (independent of genotype, all F values < 1). The least abundant protein was STX1A with an average level of 2.34±0.62ng/mg (Fig.5I) and, as with the other two synaptic proteins, the levels of STX1A were comparable between all cohorts (F values < 1).

**Figure 5:**
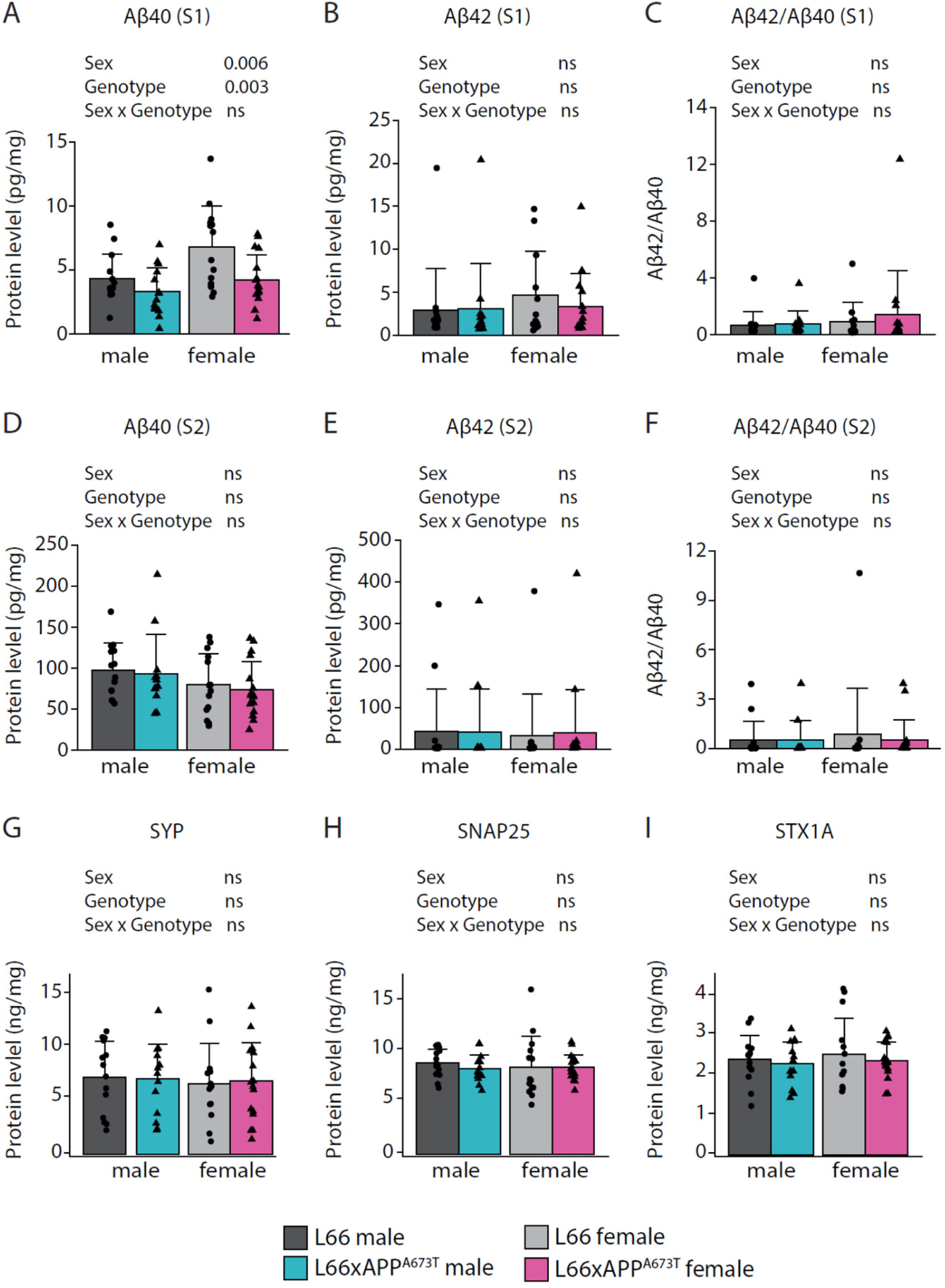
Quantification of Aβ and synaptic proteins in L66 and L66xAPP^A673T^ male and female mice. Mouse Aβ40, Aβ42 and their Aβ42/Aβ40 ratio were quantified in RIPA-soluble (A-C) and insoluble fractions (D-E) prepared from whole brain homogenates. (G) SYP, (H) SNAP25, (I) and STX1A were quantified in RIPA-soluble S1 fractions. Raw data for Aβ and synaptic proteins were normalised to protein levels and are shown as individual values, with group mean and S.D. Data were analysed using 2-way ANOVA with sex and genotype as independent variables. Aβ data was square root (A-D), or Box-Cox (E, F) transformed for analysis. Synaptic protein data (G-I) did not require transformation. Where samples could not be included due to low sample volume (Aβ quantification) this is indicated in sample sizes in brackets below. L66: males=14 (S1 Aβ42:13), females =14; L66xAPP^A673T^: males=14 (S2 Aβ40:12, S2 Aβ42:13), females=17. Abbreviations: ns: not significant, SNAP25: synaptosomal associated protein 25kDa, STX1A: syntaxin 1A, SYP: synaptophysin.

Pearson correlation matrices were generated to further investigate any differences between genotypes in terms of correlations between tau and Aβ pathology and synaptic protein markers. In male mice there was a difference between L66 and L66xAPP^A673T^ correlation matrices both visually and statistically (Fig. S1A and S1B, *p* < 0.001, see supporting information). Similarly, correlation matrices of female L66 and L66xAPP^A673T^ were also significantly different (Fig. S1C and S1D, *p* < 0.001, see supporting information). A direct comparison in terms of amyloid and synaptic markers with aggregated and non-pTau levels, however, can be obtained from correlation matrices comparing males and female mice of the different genotypes (Fig. 6A-D). While in L66 males non-pTau and Aβ levels in S1 showed negative correlations, especially with Aβ42, this correlation was lost or reduced in L66xAPP^A673T^ males (Fig. 6A). Meanwhile, aggregated tau levels showed slight positive correlations with Aβ in S1 which was not the case in L66xAPP^A673T^ crosses (Fig. 6B). On the other hand, correlations between Aβ42 and aggregate tau (in S2) were much stronger in L66xAPP^A673T^ males than L66. Further, a strong negative correlation between aggregated tau and STX1A was seen in L66 but not L66xAPP^A673T^. In females, associations between non-pTau or aggregated tau and synaptic proteins and Aβ were generally weaker than in males (Fig.6C,D). There was a slight negative correlation between S2 Aβ42 and the Aβ42/Aβ40 ratio and non-pTau in L66 females that was reduced in L66xAPP^A673T^whereas the opposite was the case for Aβ42 and Aβ42/Aβ40 in S1 (Fig.6C). Aggregated tau showed a slight negative correlation with Aβ40 and Aβ42 in S1 in L66 females that was lost in L66xAPP^A673T^ (Fig.6D).

**Figure 6:**
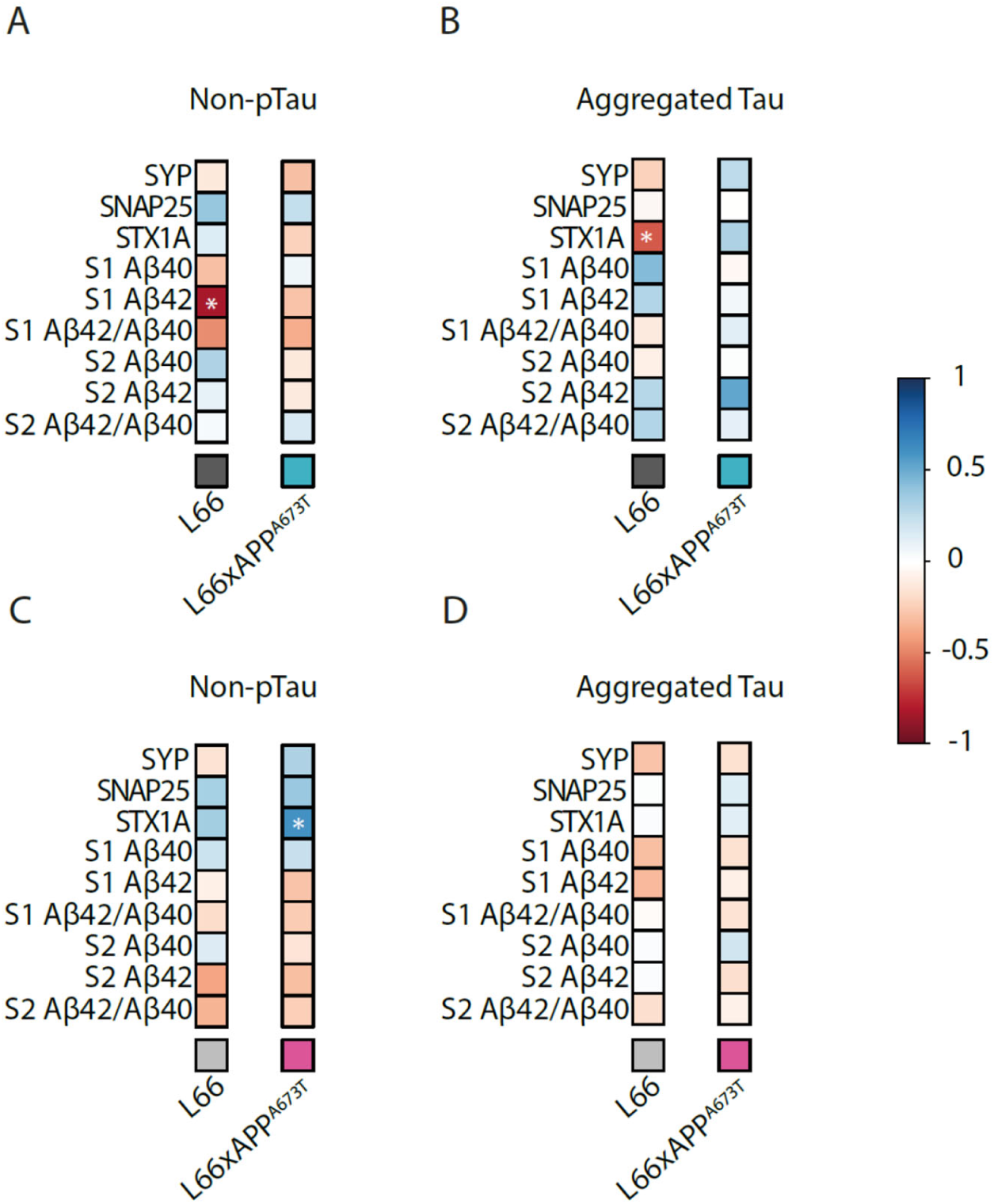
Correlation matrices between tau, Aβ and synaptic proteins. Pearson correlation matrices between non-pTau and aggregated tau with the synaptic proteins SYP, SNAP25, STX1A, as well as Aβ40, A β42 and their ratio in S1 and S2 are displayed for L66 male (A), L66xA673T male (B), L66 female (C), and L66xA673T female (D) with blue for positive correlations, red for negative correlations and white where no correlation was seen (* p < 0.05). SYP, SNAP25, STX1A, non-pTau, aggregated tau, Aβ40 and A β42 were quantified in S1 and S2 brain homogenate fractions using ELISA. HT7-tau, 27/499-tau, and 7/51-tau were quantified using immunohistochemistry (averaged across brain regions). Data were analysed using Jennrich test to detect differences between matrices. Abbreviations: SNAP25: synaptosomal associated protein 25kDa, STX1A: syntaxin 1A, SYP: synaptophysin.

### Icelandic mutation and behaviour

We also examined whether any pathological differences between and L66 and L66xAPP^A673T^ mice may translate to functional consequences in terms of behaviour and thus tested these mice using six different paradigms assessing motor (Fig.7) and neuropsychiatric symptoms including social memory (Fig.8), all known to be associated with dementia. Tests were performed in the sequence as described here.

**Figure 7:**
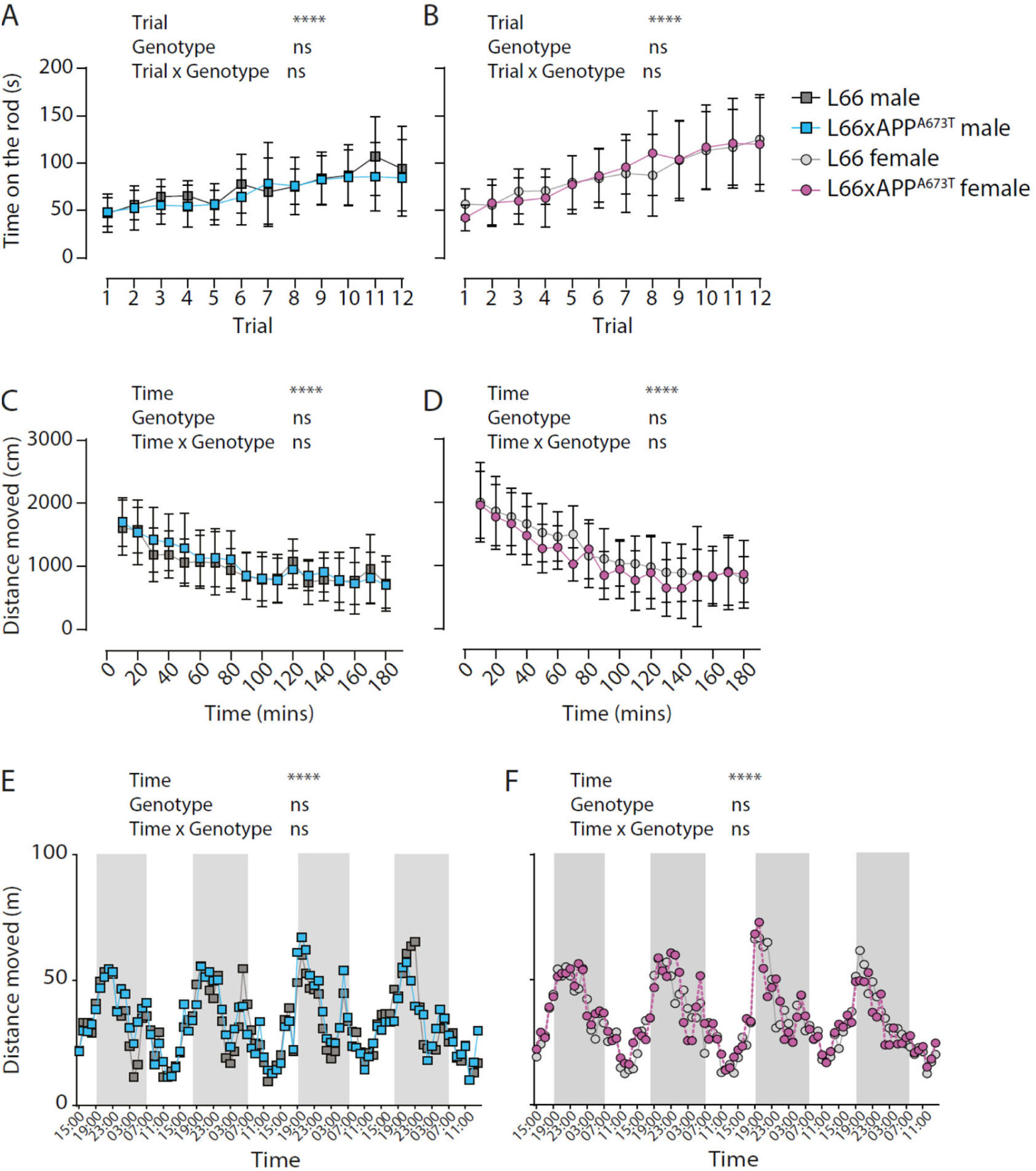
Behaviour on the rotarod and in the PhenoTyper of L66 and L66xAPP^A673T^ mice. Motor function was assessed using a four-lane rotarod system was used. Each mouse was given a total of 12 trials (T1-T12) over three days and the time spent on the rod was plotted for male (A) and female (B) mice. Activity in the PhenoTyper during the first 180 minutes (habituation) was assessed as distance moved for male (C) and female (D) mice. Additionally, the circadian activity (distance moved) over weekdays was quantified in hourly bins in the PhenoTyper for male (E) and female (F) mice and the active (dark) phase from 7 pm to 7 am is highlighted in grey. No differences between genotypes were observed between genotypes in any metric. Data are shown as either group mean and S.D. (A-D), or as group mean only for clarity (E-F). Data were analysed using 2-way ANOVA with genotype and trial (A-B) and genotype and time (C-F) as independent variables (**** p < 0.0001). Abbreviations: ns: not significant.

**Figure 8:**
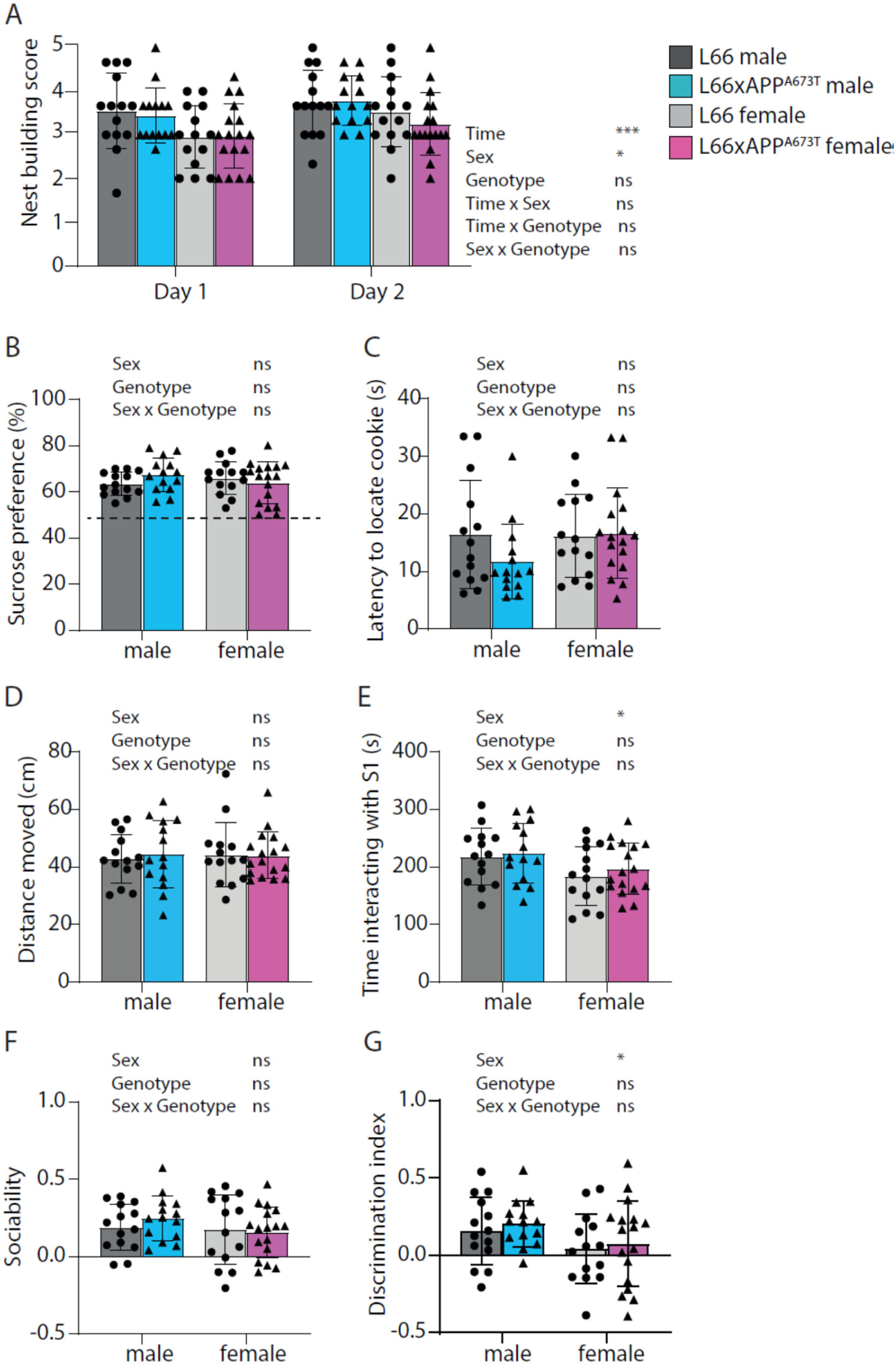
Behaviour during nestbuilding, sucrose preference, buried cookie, and social interaction tests of L66 and L66xAPP^A673T^ mice. Nestbuilding ability was accessed after a period of 16 hrs (Day 1) and 48 hrs (Day 2) following the introduction of the nestlet into home cages (A). Anhedonic-like behaviour and olfaction were assessed using the sucrose preference test (B) and the buried cookie test (C), respectively. Lastly, for the social interaction test, distance moved was determined during habituation (D), time spent interacting with the S1 stranger during the sociability phase (E), and time spent with novel stranger S2 (F), and the discrimination index (G) were analysed for the social recognition phase. No genotype related differences were recorded. Data are shown as individual values, with group mean and S.D., and were analysed using repeated measures ART ANOVA with time, sex and genotype as independent variables (A, * p < 0.05; *** p < 0.001), or 2-way analysis of variance (ANOVA) with sex and genotype as independent variables (B-G, * p < 0.05; ** p < 0.01). Dotted line in B indicates 50% level of chance. Abbreviations: ns: not significant.

Motor competence was examined using the rotarod. Motor performance, expressed as time spent on the rotating rod was similar for L66 and L66xAPP^A673T^ male (F values < 1; Fig.7A) and female cohorts (F values < 1; Fig.7B). Similarly, motor learning seen as increased time on the rod between trials and sessions occurred in all cohorts (males F_Trial_(4.405,114.5) = 11.69; females F_Trial_(5.277, 153.0) = 25.18; *p* values < 0.0001).

Home cage activity offers the possibility for long-term monitoring and the ability to assess and track behaviour in a relatively stress-free environment, and this was done in the current study using the PhenoTyper home cage observation system (Grieco et al., 2021; Robinson et al., 2013, 2018; Robinson & Riedel, 2014). The first three hours in the PhenoTyper are considered as habituation to a novel environment and the distance moved during this time is plotted separately as a function of time (Platt et al., 2011). Male mice of both genotypes habituated similarly to the new environment, as shown by the decreased distance covered over time (F_Time_(6.046,151.1) = 17.30, *p* < 0.0001; Fig.7C), and there were no significant differences between L66 and L66xAPP^A673T^ males (F_Genotype_(1, 25) = 0.32, *p* = 0.58). Female mice also showed similar habituation over time (F_Time_(5.907, 159.5) = 30.07, *p* < 0.0001; Fig.7D), and, again, no differences between genotypes were observed (F_Genotype_(1, 27) = 1.31, *p* = 0.26). Long-term monitoring of the average distance travelled in a 24hr period over the 4 days of home cage observation also did not yield significant differences between L66 and L66xAPP^A673T^ males (F_Genotype_(1, 2304) = 1.16, *p* = 0.28; Fig.7E) or females (F_Genotype_(1, 2588) = 0.50, *p* = 0.48; Fig.7F). All cohorts showed typical circadian rhythms with heightened peak ambulatory activity during the dark phase (7 p.m. until 7 a.m. highlighted in grey) and are more sleep prone during the light phase (males F_Time_(95, 2304) = 9.48; females F_Time_(95, 2588) = 12.62; *p* values < 0.0001).

The neuropsychiatric assessment included nestbuilding, sucrose preference, buried cookie, and social interaction testing (Fig. 8).

Apathy-like behaviour was measured using the nestbuilding test (Robinson et al., 2024). Readings were taken at 24 and 48 hours after the introduction of nestlets using a five-point scale. After 24 hours, all mice achieved a score between 2 and 5, with a score of 1 indicating absence of any nestlet shredding and a score of 5 indicating complete shredding and perfect nest construction. No significant differences were observed between genotypes (Fig. 8A, F_Genotype_<1), but male mice achieved significantly higher scores than female mice (F_Sex_(1, 55) = 6.06, *p* = 0.017). Additionally, all cohorts improved their nestbuilding abilities at 48 compared to 24 hours (F_Time_(1, 55) = 13.48, *p* = 0.0005). A lack of differences in anhedonia between L66 and L66xAPP^A673T^ mice was further confirmed using the sucrose preference test (Robinson et al., 2024). Data of fluid consumption revealed that all groups preferred sucrose over water with one-sample t-tests confirming that the preference for L66 and L66xAPP^A673T^ crosses were significantly above the 50% level of chance (Fig. 8B, all *p* values < 0.0001). No overall genotype or sex differences were observed for either sucrose preference (F values < 1) or total fluid intake (data not shown). The buried cookie test was used to assess olfactory ability and motivation (Fig. 8C). As no overall differences were observed between the readouts from manually timed assessments and the automated tracking software, they both were deemed to be equally reliable. Therefore, the automated testing was used for the purpose of this analysis. No genotype or sex differences were observed in the mean latency to locate the cookie during the two buried cookie trials (trials 2 and 3) with all groups locating the cookie within an average of 11-17 seconds (F values < 1). Further analysis confirmed sex differences with the L66xAPP^A673T^ males presenting with heightened activity compared to the females (*p* = 0.0017; data not shown). The results from social interaction and recognition testing are summarised in Fig. 8D-G. During the habituation phase all mice presented with similar levels of locomotor activity with no overall differences observed between genotypes or sexes (F values < 1, see Fig. 8D). Similarly, all groups spent a considerable amount of time adjacent to the cylinder containing the stranger mouse S1 during the sociability phase (Fig. 8E). This amount of time was considerably higher than the time spent with the empty cylinder (main effect of zone: (F(1,55) = 72.62, *p* < 0.0001) and this is reflected in the discrimination index for the sociability phase (Fig. 8F). While there was no main effect of sex, genotype or interaction (F’s < 1.3), all groups displayed a significant bias the interaction zone of S1 (one sample t-test, all p’s < 0.013).

For social memory, the time bias for stranger S2 was above chance only for the males, but not the females (t’s > 2.5, p’s < 0.02 for males, t’s < 1.2, ns for females). Globally, there were no differences observed between L66 and L66APP^A673T^ animals for the discrimination index (F < 1) and only a main effect of sex occurred (F(1,55) = 4.2, p = 0.04). time spent interacting with the S1 mouse (Fig. 8E). During the social recognition test, all genotypes/sexes displayed a preference for the novel stranger mouse S2, spending an increased amount of time exploring S2 compared to the familiar S1 mouse (F(1,55) = 17.28, *p* < 0.0001; data not shown) with no genotype differences for time spent with the novel S2 mouse (Fig. 8F) or the discrimination index (Fig. 8G). Sex differences were apparent during the test with both L66 and L66xAPP^A673T^ female mice displaying reduced overall exploration compared to their male counterparts (*p* values < 0.0001, data not shown). Female mice of both genotypes also spent less time interacting with the S1 mouse relative to the male mice (F(1,55) = 5.78; *p* = 0.02; Fig. 8E). Further sex differences were also present, where female mice had a lower discrimination index than male mice (F(1, 55) = 4.244; *p* = 0.0441; Fig. 8G).

## Discussion

The APP^A673T^ Icelandic mutation has been shown to reduce the risk of AD through the reduction of Aβ production (Jonsson et al., 2012; Martiskainen et al., 2017). However, its effect on tau pathology is not well studied and has never been investigated in tau transgenic mice. In this study, we have therefore examined whether the addition of the A673T mutation to the murine APP gene can directly reduce tau pathology and thereby rescue behavioural deficits previously reported for L66 tau transgenic mice (Melis et al., 2015; Robinson et al., 2024).

### A673T Aβ has a marginal impact on tau pathology and soluble Aβ40 levels in L66 mice in vivo

L66 mice overexpress the aggregation-prone mutant form P301S of the longest human tau isoform. The mice show tau accumulation in cells in various brain regions and the aggregated state for some of these tau species has been verified by binding to Bielschowsky silver and primulin (Melis et al., 2015). Furthermore, synaptic accumulation of tau in L66 mice has been confirmed by immunohistochemical co-localisation with synaptic markers and by co-purification with synaptic proteins in synaptosomal and synaptic vesicle preparations (Cranston et al., 2024; Lemke et al., 2020). Reactivity of the various tau species in L66 has been extensively investigated histopathologically using a large set of monoclonal antibodies. These include mAb 7/51 targeting an epitope within the microtubule domain, HT7 targeting an epitope within proline-rich domain of tau, and mAb 27/499 targeting its N-terminal domain (Lemke et al., 2020). These three antibodies have been applied in the current study to enable a thorough examination of the abundance of distinct tau species in different regions of the L66 brain (Lemke et al., 2020; Melis et al., 2015). However, here we report only minimal differences in tau species in L66xAPP^A673T^ crosses compared with L66 tau mice. Although we found a reduction of 7/51-reactive tau in PFC in the range of 25 and 38% (in males and females, respectively), no further differences between genotypes for other brain regions of interest and antibodies emerged. This is surprising given that the three antibodies, 7/51, HT7 and 27/499 span a number of domains across the tau molecule and L66 mice showing intense immuno-positive labelling that is absent from non-transgenic wild-type control mice (Lemke et al., 2020).We take this as a lack of efficacy of the APP^A673T^ mutation to mechanistically counter the overexpression of tau in 6-month-old L66 mice. This was confirmed for soluble and insoluble fractions in which no overt differences in tau levels were observed between genotypes. The ELISAs used to measure such changes were highly specific for L66 tau relative to non-transgenic wild-type mice (unpublished data).

Previous work has suggested that the APP^A673T^ Icelandic mutation can protect against Aβ pathology in Aβ-transgenic mice und Aβ-knock-in rats (Shimohama et al., 2024; Tambini et al., 2020). In our model, in which the mutation was introduced into the murine APP gene and the resultant mice crossed with L66, soluble mouse Aβ40 was reduced by 23 to 38% (in males and females, respectively) and this reduction of Aβ40 is in line with the reduction seen for mAb 7/51-reactive tau in PFC. Intriguingly, a recent study used APPswe/PS1dE9 transgenic mice and inoculated them with either recombinant non-mutant human Aβ or human Aβ containing the A673T mutation. No changes in Aβ levels but a decrease in phospho-tau pathology in the A673T-Aβ-treated group were observed and this remains unexplained (Célestine et al., 2024). Here, we did not measure phospho-tau in L66 brains because the aggregation of non-phosphorylated tau is correlated with the onset of cognitive impairment in mice (Melis et al., 2015), and the phosphorylation of tau inhibits tau-tau binding and is preceded by aggregation of non-phosphorylated tau (Lai et al., 2016; Schneider et al., 1999). These latter data strongly suggest a prominent role of phosphorylation-independent processing of tau in both its aggregation and associated decline in cognition. A recent exploratory study in 6 non-AD patients (unconfirmed idiopathic normal pressure hydrocephalus cases) comparing CSF of three APP^A673T^ carriers to three age- (and sex) matched control subjects reported that disease relevant soluble APP-β and Aβ42 levels were significantly reduced in CSF of APP^A673T^ carries. Yet, soluble APP-α, Aβ40, total tau and phosphorylated tau (p-tau 181) were not altered (Wittrahm et al., 2023), questioning the efficacy of the ‘protective mutation’ in terms of tau expression. While this is a small study cohort and needs verification, the work seems to suggest that detection of protective A673T amyloid levels may differ between compartments and be specific to CSF. We here did not investigate CSF/plasma tau in the current study but reasoned that detection of disease-relevant markers directly at source, i.e. neurones and within brain may be a more appropriate means for differentiation of phenotypes. Moreover, recent work in L66 has shown a highly significant correlation between insoluble brain tau and p-tau 217 plasma tau (Penny LK, 2025). Consequently, CSF tau in L66 would reflect brain tau and is unlikely to be reduced following the introduction of the APP^A673T^ mutation in our L66 tau transgenic line. Interestingly, the A673T mutation also altered correlations between the Aβ and both non-pTau and aggregated tau. This would suggest that APP^A673T^ has unique effects on tau pathology that differ from wildtype Aβ which is interesting considering that both proteins can act in complex synergy (Busche & Hyman, 2020). Since a reduction of endogenous murine APP exacerbated tau pathology in an APP-knock out and tau overexpressing mouse model (Tg30) (Vanden Dries et al., 2017), it is not inconceivable that small reductions of Aβ40/42 as seen in our L66xAPPA673T crosses does not reduce tau production and is unable to rescue either tau pathology or the behavioural responses of L66 mice.

### A673T Aβ does not modify synaptic protein expression in L66 in vivo

Synaptic dysfunction that contributes to memory impairment and synapse loss, confirmed by post-mortem analyses of AD brain tissue, is the strongest correlate for cognitive decline (Anschuetz et al., 2024; De Wilde et al., 2016; Spires-Jones, 2014). The presynaptic proteins SYP and SNAP25 were chosen as established markers for synapse loss in AD, as well as in tau- and Aβ-based mouse models (Anschuetz et al., 2024, 2025; De Wilde et al., 2016). Our previous work established that tau aggregation induces alterations in synaptic proteins and synaptic transmission in L66 mice (Schwab et al., 2021). This made it reasonable to confirm whether the protective mutation in the APP gene could correct the abnormal levels of synaptic proteins. Six-month old L66 mice are particularly sensitive to reductions in structural and release-competent presynaptic proteins such as SNAP25 and STX1A, and reduction of tau using a small molecule aggregation inhibitor resulted in partial recovery of normal synaptic protein expression (Schwab et al., 2024; Schwab et al., 2025). In the current study, APP^A673T^ did not seem to modulate the expression of the three investigated presynaptic proteins SYP, SNAP25 and STX1A. This is likely the result of the lack of substantial modulation of tau and/or Aβ in the L66xAPP^A673T^ crosses. However, there was a strong negative correlation between levels of aggregated tau and STX1A in L66 male mice that was absent in L66xAPP^A673T^. In female L66xAPP^A673T^, there was an enhanced positive correlation between STX1A and non-pTau compared to L66. The lack of negative association between STX1A and aggregate tau could suggest a small protective effect of L66xAPP^A673T^, making synapses less sensitive to synaptic protein loss in response to tau aggregation. No such association was seen for either genotype in females, suggesting there may also be a sex-specific component to this.

### A673T Aβ does not modulate motor and neuropsychiatric phenotypes in L66 tau transgenic mice

The APP^A673T^ mutation was also associated with increased cognitive performance in carriers without AD compared to noncarriers based on the Cognitive Performance Scale, suggesting a possible protective effect of the Icelandic mutation that may be unrelated to AD and/or Aβ/tau pathologies (Jonsson et al., 2012). This notion was based on a case study of a cognitively normal centenarian with vascular amyloid pathology and Braak stage 3 tau pathology (Kero et al., 2013). We therefore reasoned that our L66xAPP^A673T^ crosses could benefit from this mutation independent of dementia-related pathological benefits and performed a behavioural phenotyping study despite the lack of any substantial tau and/or Aβ reduction. Using six different behavioural tests addressing motor function, olfaction, depression and apathy-like behaviour, as well as motivation, exploration and sociability/social memory, we failed to find any overt differences between L66 and L66xAPP^A673T^ crosses. While not in line with our hypothesis, it needs to be considered that only very few studies have investigated the effect of the Icelandic mutation on behaviour in mice and results are inconsistent. For example, inoculation of A673T Aβ into Aβ-transgenic mice rescued spatial memory deficits in the Morris water maze via reduction of tau but not Aβ pathology (Célestine et al., 2024). A related study investigated the APP^A673V^ mutation, which has previously been described to reduce Aβ if expressed in a heterozygous background (Cantu et al., 2017; Di Fede et al., 2009). After induction of a traumatic brain injury in the 3xTg-AD model, mice were inoculated with vehicle or APP^A673V^. Improved locomotor activity in the open field and cognitive behaviour in the Y-maze was amongst the beneficial effects induced by the protective Aβ peptide (Diomede et al., 2023). Discrepancies in behavioural outcomes are most likely related to the use of distinct AD models (APPswe/PS1dE9 (Célestine et al., 2024), 3xTg-AD (Diomede et al., 2023) or L66 in the current study) but also to the different approach for APP^A673T^ delivery (inoculation vs genetic expression). No systematic comparison between the approaches is yet available but urgently needed. Moreover, earlier work revealed that the accumulation of tau in L66 lead to sensorimotor/motor deficiencies and neuropsychiatric symptoms e.g. using rotarod, sucrose preference and nestbuilding paradigms at 6 months of age, but not a cognitive phenotype (Melis et al., 2015; Robinson et al., 2024). It is therefore conceivable that benefits exerted by APP^A673T^ are specific to memory functions not examined here, yet which requires further study.

In summary, we here show that the APP^A673T^ Icelandic mutation does not modulate motor and neuropsychiatric behaviours in L66 tau transgenic mice, possibly due to the lack of modification of tau pathology in 6-month-old L66xAPP^A673T^ mice.

## INSTITUTIONAL REVIEW BOARD STAT EMENT

All animal experiments were performed in accordance with the European Communities Council Directive (63/2010/EU) and a project licence with local ethical approval under the UK Animals (Scientific Procedures) Act (1986) and its Amended Regulations (2012) and comply with the ARRIVE 2.0 guidelines. No human samples were used in this study.

## LIST OF ABBREVIATIONS

AD: Alzheimer’s disease
APP: amyloid precursor protein
ART: Aligned Rank Transform
Aβ: amyloid β-protein
BCA: bicinchoninic acid
CA1: cornu ammonis
CB: cerebellum
CTX: visual cortex
DG: dentate gyrus
IHC: immunohistochemistry
L66: line 66 tau transgenic mice
mAb: monoclonal antibody
non-pTau: non-phosphorylated tau
PFC: prefrontal cortex
S.D.: standard deviation
SNAP25: synaptosome associated protein 25kDa
STX1A: syntaxin 1A
SYP: synaptophysin

## ACKNOWLEDGEMENTS

The authors acknowledge Douglas Eydmann for support with immunohistochemistry.

## AVAILABILITY OF DATA AND MATERIALS

Data will be made available on reasonable request.

## AUTHORS’ CONTRIBUTIONS

Anne Anschuetz: Investigation, Data curation, Software, Formal analysis, Visualization, Writing - Original Draft; Lianne Robinson: Conceptualization, Data Curation, Formal analysis, Writing - Review & Editing; Miguel Mondesir: Investigation, Data curation; Valeria Melis: Resources; Bettina Platt: Conceptualization, Resources, Writing - Review & Editing; Charles R. Harrington: Funding acquisition, Writing - Review & Editing; Gernot Riedel: Conceptualization, Supervision, Project administration, Funding acquisition, Writing - Review & Editing; Karima Schwab: Conceptualization, Supervision, Investigation, Data curation, Formal analysis, Visualization, Writing - Original Draft.

---

**Figure S1:**
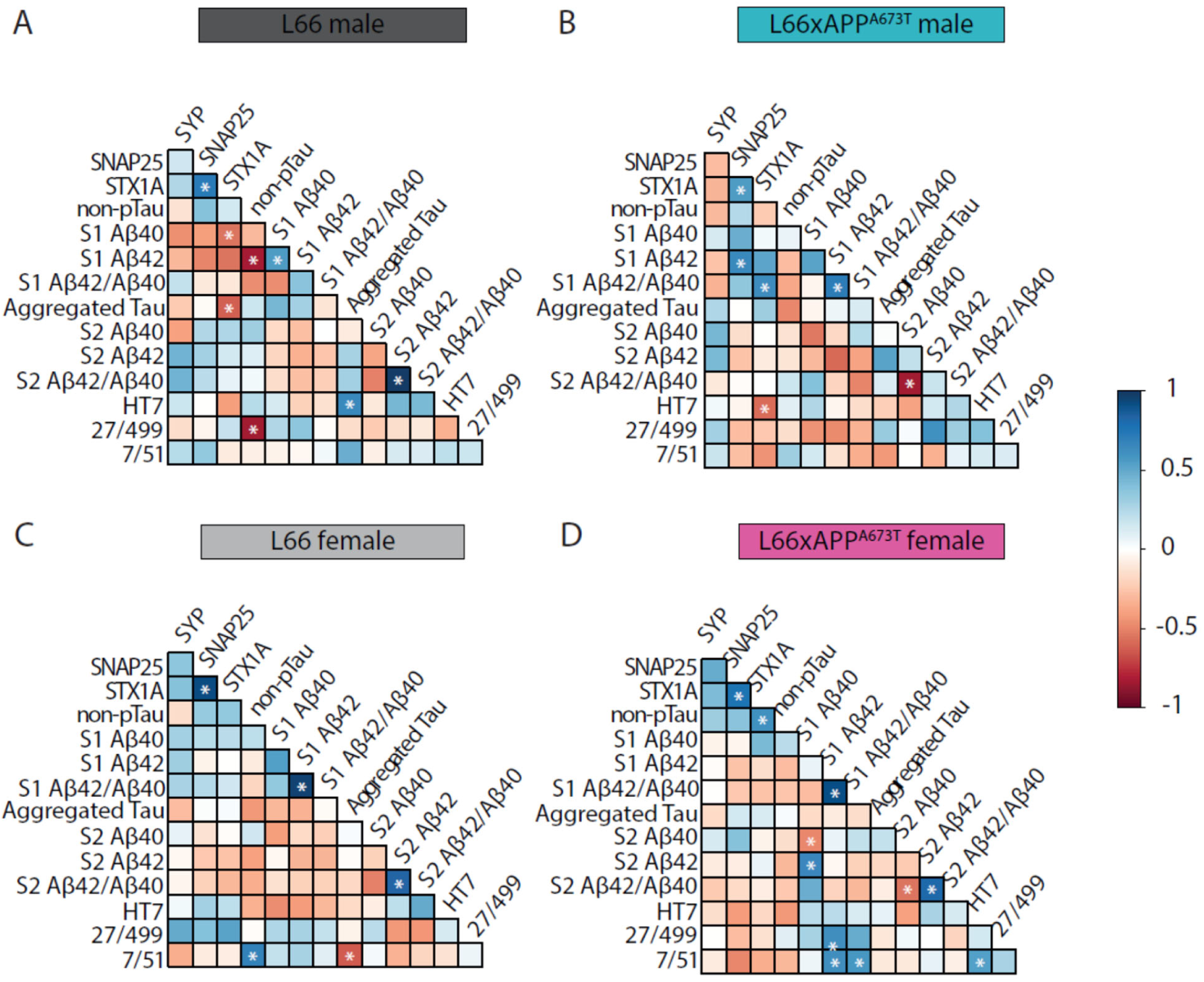
Correlation matrices between tau, Aβ and synaptic proteins of L66 and L66xAPP^A673T^ mice. Pearson correlation matrices between the synaptic proteins SYP, SNAP25, STX1A, as well as non-pTau, aggregated tau, Aβ40 and Aβ42 as well as their ratio are displayed for L66 male (A), L66xA673T male (B), L66 female (C), and L66xAPP^A673T^ female (D). Blue denotes positive correlations, red for negative correlations and white indicates where no correlations were seen (* p < 0.05). SYP, SNAP25, STX1A, non-pTau, aggregated tau, Aβ40 and A β42 were quantified in S1 and S2 brain homogenate fractions using ELISA. HT7-tau, 27/499-tau, and 7/51-tau were quantified using immunohistochemistry (averaged across brain regions). Data were analysed using Jennrich test to detect differences between matrices. Abbreviations: SNAP25: synaptosomal associated protein 25kDa, STX1A: syntaxin 1A, SYP: synaptophysin.

